# Learning the Drug-Target Interaction Lexicon

**DOI:** 10.1101/2022.12.06.519374

**Authors:** Rohit Singh, Samuel Sledzieski, Lenore Cowen, Bonnie Berger

## Abstract

Sequence-based prediction of drug-target interactions has the potential to accelerate drug discovery by complementing experimental screens. Such computational prediction needs to be generalizable and scalable while remaining sensitive to subtle variations in the inputs. However, current computational techniques fail to simultaneously meet these goals, often sacrificing performance on one to achieve the others. We develop a deep learning model, ConPLex, successfully leveraging the advances in pre-trained protein language models (“PLex”) and employing a novel protein-anchored contrastive co-embedding (“Con”) to outperform state-of-the-art approaches. ConPLex achieves high accuracy, broad adaptivity to unseen data, and specificity against decoy compounds. It makes predictions of binding based on the distance between learned representations, enabling predictions at the scale of massive compound libraries and the human proteome. Furthermore, ConPLex is interpretable, which enables us to visualize the drug-target lexicon and use embeddings to characterize the function of human cell-surface proteins. We anticipate ConPLex will facilitate novel drug discovery by making highly sensitive and interpretable in-silico drug screening feasible at genome scale. ConPLex is available open-source at https://github.com/samsledje/ConPLex.

**Significance Statement:** In time and money, one of the most expensive steps of the drug discovery pipeline is the experimental screening of small molecules to see which will bind to a protein target of interest. Therefore, accurate high-throughput computational prediction of drug-target interactions would unlock significant value, guiding and prioritizing promising candidates for experimental screening. We introduce ConPLex, a machine learning method for predicting drug-target binding which achieves state-of-the-art accuracy on many types of targets by using a pre-trained protein language model. The approach co-locates the proteins and the potential drug molecules in a shared feature space while learning to contrast true drugs from similar non-binding “decoy” molecules. ConPLex is extremely fast, which allows it to rapidly shortlist candidates for deeper investigation.

---

**I**n the drug discovery pipeline, a key rate-limiting step is the experimental screening of potential drug molecules against a protein target of interest. Thus, fast and accurate computational prediction of drug-target interactions (DTIs) could be extremely valuable, accelerating the drug discovery process. One important class of computational DTI methods, molecular docking, uses 3D structural representations of both the drug and target. While the recent availability of high-throughput accurate 3D protein structure prediction models (1–3) means that these methods can be employed starting only from a protein’s amino acid sequence, the computational expense of docking (4) and other structure-based approaches (e.g., rational design (5), active site modeling (6), template modeling (7, 8)) unfortunately remains prohibitive for large-scale DTI screening. An alternative class of DTI prediction methods use 3D structure only implicitly, making rapid DTI predictions when the inputs consist only of a molecular description of the drug (such as the SMILES string (9)) and the amino acid sequence of the protein target. This class of *sequence-based* DTI approaches enables scalable DTI prediction, but there have been barriers to matching the levels of accuracy obtained by structure-based approaches.

In this paper we introduce ConPLex, a new rapid purely sequence-based DTI prediction method that leverages protein language models (PLMs), and show it can produce state of the art performance on the DTI prediction task at scale. The advance provided by ConPLex comes from two main ideas that together overcome some of the limitations of previous approaches. While many methods have been proposed for the sequence-based setting of the DTI problem (10) (e.g., using secure multi-party computation (11), convolutional neural networks (12) or transformers (13)), their protein and drug representations are constructed solely from DTI ground truth data. The high level of diversity among the DTI inputs, combined with the limited availability of DTI training data, limit the accuracy of these methods and their generalizability beyond their training domain. Furthermore, the methods that do generalize often do so by sacrificing fine-grained specificity, i.e., are unable to distinguish true-positive binding compounds from false positives with similar physico-chemical properties (“decoys”).

In contrast, the “PLex” (Pre-trained Lexographic) part of ConPLex helps alleviate the problem of limited DTI training data. As we showed in our preliminary work (14), one way to get around the limited size of DTI data sets that has hampered the quality of the representations learnt by previous methods is to transfer learned proteins representations from pre-trained protein language models to the DTI prediction task. These representations encode structural insights and benefit from being trained on the much larger corpus of single protein sequences (14). Starting with the PLM models, our second insight directly addresses the fine-grained specificity problem in our architecture by using the “Con” (Contrastive learning) part: a novel, protein-anchored contrastive co-embedding that co-locates the proteins and the drugs into a shared latent-space. We show that this co-embedding enforces separation between true interacting partners and decoys to achieve both broad generalization and high specificity.

Putting these two ideas together gives us ConPLex, a novel representation learning approach that enables both broad generalization and high specificity. We show that ConPLex enables more accurate prediction of DTIs than competing methods while avoiding many of the pitfalls suffered by currently available approaches. Thus, our work constitutes a concrete demonstration of the power of a well-designed transfer learning approach that adapts foundation models for a specific task (15, 16). In particular, we found that the performance of existing sequence-based DTI prediction methods could be sensitive to variation in drug-vs-protein coverage in the data set, whereas ConPLex performs well in multiple coverage regimes. Indeed, ConPLex performs especially well in the zero-shot prediction setting where no information is available about a given protein or drug at training time.

ConPLex can also be adapted beyond the binary cases to make predictions about binding affinity. Furthermore, the shared representation also offers advantages beyond prediction accuracy. The co-embedding of both proteins and drugs in the same space offers intepretability, and we show that distances in this space meaningfully reflect protein domain structure and binding function: we leverage ConPLex representations to functionally characterize cell-surface proteins from the Surfaceome database (17), a set of 2,886 proteins localized to the external plasma membrane that participate in signaling and are likely able to be easily targeted by ligands.

ConPLex is extremely fast: as a proof-of-concept, we make predictions for the human proteome against all drugs in ChEMBL (18) (≊ 2 × 10^10^ pairs) in just under 24 hours. Thus, ConPLex has the potential to be applied for tasks which would require prohibitive amounts of computation for purely structure based approaches or less efficient sequence-based methods, such as genome-scale side-effect screens, identifying drug re-purposing candidates via massive compound libraries searches, or *in silico* deep mutational scans to predict variant effects on binding with currently approved or potential new therapeutics. We note that most DTI methods require significant computation on pairs of drugs and targets (i.e., have quadratic time-complexity). Because ConPLex predictions rely only on the distance in the shared space, predictions can be made highly efficiently once embeddings (which have linear time-complexity) are computed.

## Distinguishing between low- and high-coverage DTI prediction

We benchmark performance of ConPLex and competing methods in two different regimes, which we term low-coverage and high-coverage DTI prediction (Figure 1c). We show that ConPLex outperforms its competitors in both settings, but note that separating the two regimes helps clarify an often-seen issue in the field: methods whose performance varies substantially across different proposed DTI benchmarks. Several prior attempts have been made to standardize DTI benchmarking and develop a consistent framework for model evaluation (19, 20). However, much of this work has overlooked a key aspect of benchmarking that we find to significantly affect model performance—differing per-biomolecule data coverage. We define coverage as the average proportion of drugs or targets for which a data point exists in that data set, whether that is a positive or negative interaction (**Methods**). Depending on the per-biomolecule data coverage of the benchmark data set, we claim that these benchmarks are looking at very different problems. In particular, low-coverage data sets (Figure 1a) tend to measure the broad strokes of the DTI landscape, containing a highly diverse set of drugs and targets. Such data sets can present a modeling challenge due to the diverse nature of targets covered, but allow for a broad assessment of compatibility between classes of compounds and proteins. High-coverage data sets (Figure 1b) represent the opposite trade-off: they contain limited diversity in drug or target type, but report a dense set of potential pairwise interactions. Thus, they capture the fine-grained details of a specific sub-class of drug-target binding and enable distinguishing between similar biomolecules in a particular context.

**Fig. 1.**
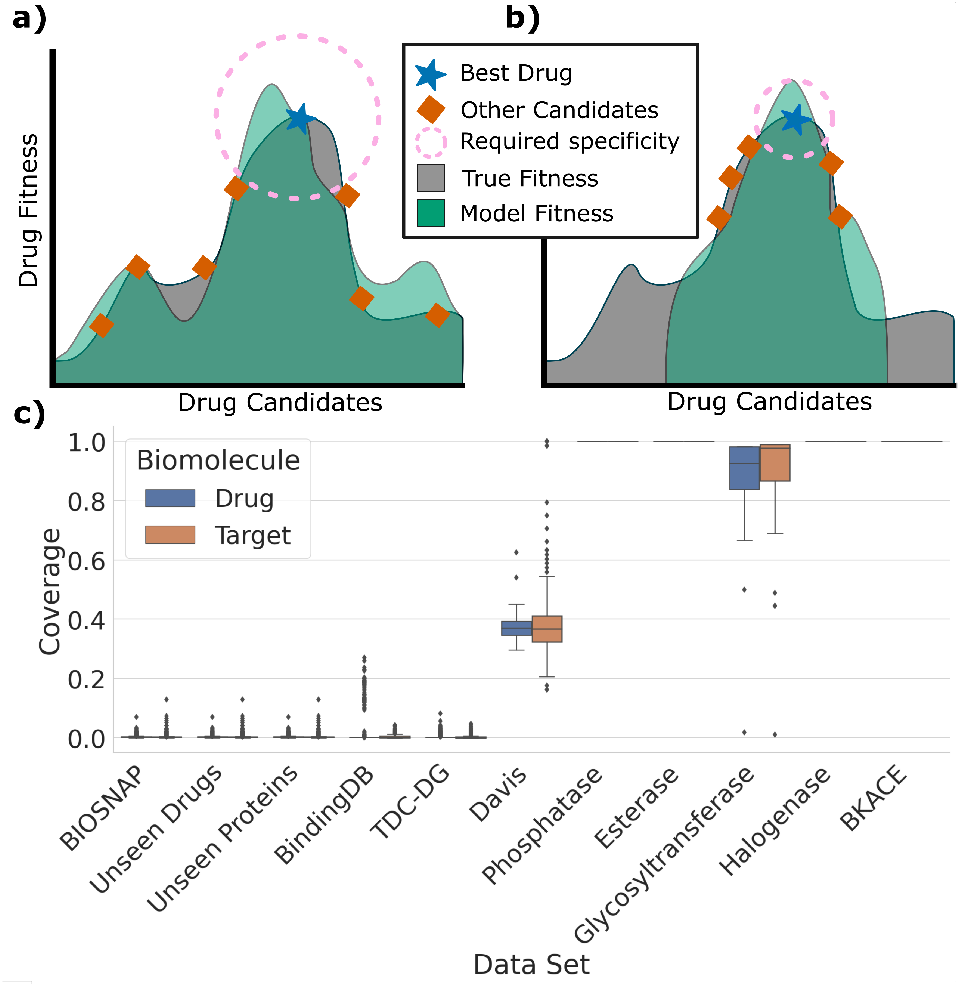
Drug-target interaction benchmarks display highly variable levels of coverage. Coverage is defined as the proportion of drugs or targets for which a data point (positive or negative) exists in that data set. High- vs. low-coverage benchmarks tend to reward different types of model performance. **(a)** In this cartoon example of a low coverage data set, drug candidates cover the full diversity of the space, and no two drugs are highly similar. A successful model does not need to be highly specific, but must accurately model the entire fitness landscape to generalize to all candidates. **(b)** For low-coverage data sets, drugs tend to be much more similar. A successful model does not need to generalize nearly as widely, but must achieve high specificity to differentiate between similar drugs. **(c)** In a review of existing popular DTI benchmark data sets, we find widely varying coverage, from data sets with nearly zero coverage (each drug/target is represented only a few times) to nearly full coverage (all drug-by-target pairs are known in the data).

**Fig. 2.**
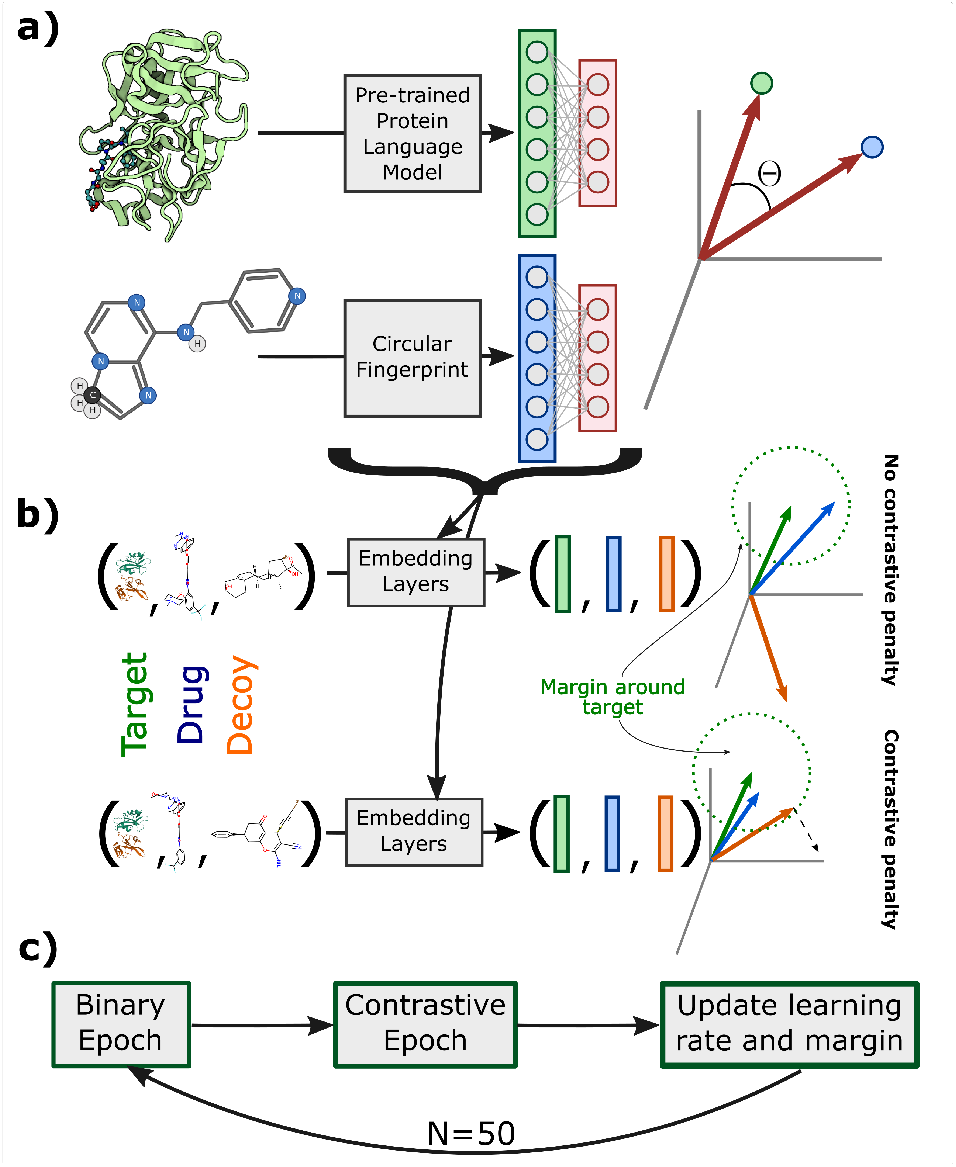
Outline of the ConPLex model architecture and training framework. ConPLex is trained in two phases, to optimize both generalizability and specificity. **(a)** Protein features are generated using a pre-trained protein language model (here ProtBert (27)), and drug features are generated using the Morgan fingerprint (26). These features are transformed into a shared latent space by a learned non-linear projection. The prediction of interaction is based on the cosine distance in this space, and the parameters of the transformation are updated using the binary cross-entropy on a low-coverage data set. **(b)** In the contrastive phase, triplets of a target, drug, and decoy are transformed in the same way into the shared space. Here, the transformation is treated as a metric learning problem. Parameters are updated using the triplet distance loss on a high-coverage data set to minimize the target-drug distance while maximizing the target-decoy distance. No additional penalty is applied if the target-decoy distance is greater than the target-drug distance plus some margin. **(c)** ConPLex is trained in alternating epochs of the binary and contrastive phase to simultaneously optimize both objectives. After each round, learning rates and the contrastive margin are updated according to an annealing scheme.

The two coverage regimes correspond to different usage cases. The low-coverage regime is relevant when applying DTI models for large-scale scans to predict interactions for a potential target against a large compound library (e.g. for drug re-purposing as in Dönertaş et al. (21) and Morselli et al. (22)), or for scanning a candidate drug against an entire proteome to identify potential adverse and off-target effects (as in Huang et al. (23, 24)). Data at this scale is often low coverage, with only a small number of known interactions for each unique biomolecule. Thus, it is important that DTI models used for these tasks are broadly applicable and can accurately generalize to many different families of proteins and drugs. However, this generalization often comes at the cost of specificity, resulting in models that are unable to distinguish between highly similar drugs or proteins.

The high-coverage regime is relevant when optimizing a particular interaction. Here, models can be trained to be highly specific to a protein family or class of drugs, so much so that a per-drug or per-target model is trained to capture the precise binding dynamics of that biomolecule (25). While such models can be effective for lead optimization, they require high coverage on the biomolecule of interest to make accurate predictions; this may not always be available. Additionally, such models lack the capacity to generalize beyond the training domain and thus cannot be used for genome- or drug bank-scale prediction.

The PLM approach of ConPLex enables strong performance in both regimes. In the low coverage regime, the strength is coming mostly from the “PLex” part, where it can leverage the effective generalization of language models to achieve state-of-the-art performance. On high-coverage data sets, the “Con” part also becomes important, since it becomes feasible to train drug- or target-specific models with high accuracy, and such models often outperform more generic models. We find that while single-task models do perform well given available data, ConPLex is able to achieve extremely high specificity in low-diversity, high-coverage scenarios, while remaining broadly applicable to protein targets with limited data. Thus, ConPLex is applicable for both large-scale compound or target screens and fine-grained, highly specific binding prediction. We discuss the issue of matching the right model to the problem domain with respect to coverage further in the **Discussion**.

## Results

### Model Overview

To achieve both generalizability and specificity, ConPLex leverages advances in both protein language modeling and metric learning. We start with pre-trained representations and learn a non-linear projection of these representations to a shared space 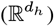. We guide the learning by alternating between two objectives over multiple iterations: a coarse-grained objective of accurately classifying DTIs, and a fine-grained objective of distinguishing decoys from drugs. The coarse-grained objective is evaluated over a low-coverage data set, which trains the model to distinguish between broad classes of drug and target, and makes initial predictions in the right “neighborhood” of the DTI space. The fine-grained objective is evaluated over a high-coverage data set, which fine-tunes the model to distinguish between true and false positive interactions in the same “neighborhood” and achieve high specificity within a class.

To featurize the inputs, here we use the Morgan fingerprint (26) for small molecules, and embeddings from a pre-trained ProtBert model (27) for proteins. We investigate other choices for features, including several other foundation PLMs in Supplementary S2. We note that our framework is flexible to different methods of featurization, and make recommendations on the selection of informative representations in the **Discussion**.

### ConPLex achieves state-of-the-art performance on low–coverage and zero-shot interactions

A key advance of Con-PLex is the use of pre-trained protein language models (PLMs) for protein representation. As foreshadowed by Scaiewicz and Levitt (29), PLMs have repeatedly been shown to encode evolutionary and structural information (30–32), and to enable broad generalization in low-coverage scenarios (33, 34). Here, we show that ConPLex achieves state-of-the-art performance on three low-coverage benchmark data sets – **BIOSNAP, BindingDB**, and **DAVIS** – where it is important to learn the broad strokes of the DTI landscape. In Table 1 we show the average area under the precision-recall curve (AUPR) over 3 random initializations of each model evaluated on a held-out test set (**Methods**). Here, we compare with several methods which use non-PLM protein features: MolTrans (13), GNN-CPI (28), and DeepConv-DTI (12). In addition, we compare to the EnzPred-CPI model from Goldman et al. (25) (developed simultaneously and independently), which uses a PLM for protein featurization but does not perform a co-embedding or utilize a contrastive training step. Finally, we compare with the single-task Ridge regression model described in (25), which trains a different model per-drug rather than a single model for the entire benchmark.

**Table 1.** ConPLex is highly accurate and generalizes broadly in low coverage settings. ConPLex outperforms several state-of-the-art methods, including EnzPred-CPI (25), MolTrans (13), GNN-CPI (28), and DeepConv-DTI (12), as well as a simple single-target Ridge regression model, on several low- and zero-coverage benchmark data sets. We report the average and standard deviation of the area under the precision-recall curve (AUPR) for 5 random initializations of each model. Metrics for models with † are taken from (13). Ridge regression cannot be applied for the Unseen Drugs data set, since a separate model is trained for each drug in the training set.

| Data Set | ConPLex | EnzPred-CPI | MolTrans | GNN-CPI† | DeepConv-DTI† | Ridge |
| --- | --- | --- | --- | --- | --- | --- |
| BIOSNAP | **0.897 ± 0.001** | 0.866 ± 0.003 | 0.885 ± 0.005 | 0.890 ± 0.004 | 0.889 ± 0.005 | 0.641 ± 0.000 |
| BindingDB | **0.628 ± 0.012** | 0.602 ± 0.006 | 0.598 ± 0.013 | 0.578 ± 0.015 | 0.611 ± 0.015 | 0.516 ± 0.000 |
| DAVIS | **0.458 ± 0.016** | 0.277 ± 0.009 | 0.335 ± 0.017 | 0.269 ± 0.020 | 0.299 ± 0.039 | 0.320 ± 0.000 |
| Unseen Drugs | **0.874 ± 0.002** | 0.844 ± 0.005 | 0.863 ± 0.005 | - | 0.847 ± 0.009 | N/A |
| Unseen Targets | **0.842 ± 0.006** | 0.795 ± 0.004 | 0.668 ± 0.045 | - | 0.766 ± 0.022 | 0.617 ± 0.000 |

Observing the strength of ConPLex to generalize on low-coverage data, we sought to evaluate its performance on fully zero-shot prediction. **Unseen drugs** and **Unseen targets** are variants of the BIOSNAP data set where drugs/targets in the test set do not appear in any interactions in the training set (**Methods**). Note that for the unseen drugs setting, the Ridge model cannot be applied since a different model must be trained for each drug that appears in the training set. We show that ConPLex achieves the best zero-shot prediction performance (Table 1), further demonstrating the applicability of the model to large-scale, very low-coverage prediction tasks.

### Contrastive learning enables high-specificity DTI mapping

Another key advance of our method is the use of contrastive learning to fine-tune model predictions on high-coverage data to achieve high specificity. Recently, Heinzinger et al. (36) demonstrated the use of semi-supervised contrastive learning for effective protein embedding-based annotation transfer. Here, we adapt contrastive learning to a fully-supervised setting and demonstrate that the contrastive training is essential to achieving specificity using DTI pairs from the Database of Useful Decoys (**DUD-E**) (35). The DUD-E data set contains 57 protein targets and drugs which are known to interact with each target. However, it also contains 50 negative “decoy” small molecules for each drug, which have similar physicochemical properties to the truly interacting small molecule, but are known to not bind the target. Thus, accurate prediction on DUD-E requires a model to achieve high-specificity and to accurately differentiate between highly similar compounds. Additionally, DUD-E contains four different classes of targets (GPCRs, kinases, proteases, and nucleases), so models must generalize across target classes (note that single task models don’t have this generalization requirement, since a different model is trained per-target).

We derive evaluation sets from DUD-E by holding out 50% of proteins in each target class for testing and using the remaining targets for training (full splits are specified in Supplementary S1). Here, we evaluate a ConPLex model trained on BIOSNAP, both with and without contrastive training on DUD-E, and show that contrastive training is essential to achieving specificity on decoys.

For each target in the DUD-E test set, we use t-SNE to visualize the target alongside all drugs and decoys using embeddings learned by both versions of the model. Figure 3a,b shows one such example, the tyrosine kinase *VGFR2*. We also show the distribution of distances in the latent space between the target embedding and the embeddings of the drugs and decoys for each model (Figure 3c, d) (*p*-values from one-sided t-test). Without contrastive training, drugs are interspersed with decoys and are far away in space from the target, while ConPLex clusters most true drugs very close to both each other and the *VFGR2* embedding.

**Fig. 3.**
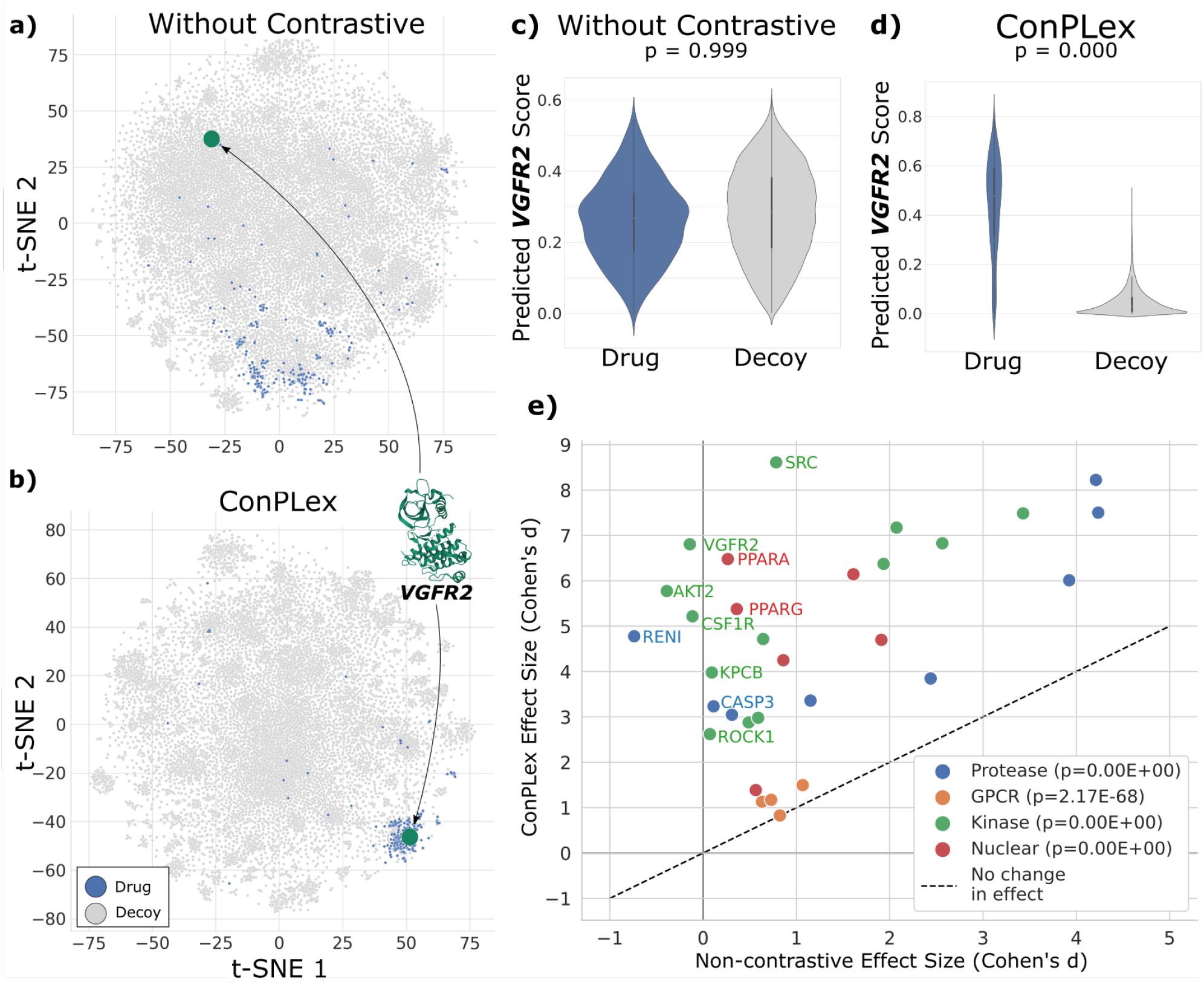
Contrastive training enables high-specificity in discriminating drugs from decoys. We demonstrate that contrastive learning is essential for ConPLex to achieve high specificity using the DUD-E (35) data set of drugs and decoys (non-binding small molecules with similar physicochemical properties to the true drugs). **(a, b)** Using t-SNE, we show the learned ConPLex latent space for *VGFR2* (green) and known drugs (blue) and decoys (grey). Without contrastive training, drugs and decoys representations do not separate and true drugs are far from their target. With contrastive training, *VGFR2* and drugs cluster very tightly compared to decoys. **(c, d)** ConPLex predictions significantly differentiate between drugs and decoys after contrastive training (*p* = 0.000 paired t-test), but do not differ at all without such training (*p* = 0.999). **(e)** We compute the effect size between drug and decoy predictions using Cohen’s *d* for all 31 targets in the test set. Targets are classified as proteases (blue), GPCRs (orange), kinases (green), and nuclear proteins (red). This effect is computed for ConPLex both with and without contrastive training. Contrastive training increases the effect size for every target (median 0.730 vs 4.716). For each class, we report the median *p*^−^value for ConPLex drug vs. decoy predictions. ConPLex performs particularly well for kinases and nuclear proteins, and more poorly for GPCRs.

In Figure 3e, we show a quantitative analysis of all 31 test-set targets. We compute the effect size (Cohen’s *d*) of the difference between predicted drug and decoy scores. We plot these effect sizes for ConPLex trained with and without contrastive training. An increase in the effect size indicates that the co-embedding distances learned by the model better represent binding specificity. The effect size increases for every target, and the median effect size between predicted true and decoy compound scores was 0.730 prior to contrastive training compared to 4.716 after. For each class of targets, we also report the median *p*-value (one-sided t-test) between drug and decoy scores predicted by ConPLex. While contrastive training has an extremely large impact on specificity in high-coverage domains, we also show that this additional training does not significantly decrease the model performance on low-coverage benchmarks via an ablation study in Supplementary S3.

In addition to evaluation on DUD-E, we also evaluate ConPLex on 5 benchmark data sets derived from family-specific enzyme-substrate screens (**Methods**). These data sets are extremely high coverage, generally including data points for all possible pairs of drugs and targets. We find that in this regime, ConPLex and other PLM based models like EnzPred-CPI have strong but highly variable performance, and are still generally outperformed by a Ridge regression model (Supplementary S5) as shown previously in (25). However, a fine scale single-task model is limited in its generalizability beyond the enzyme family on which it was trained (**Discussion**).

### Interpreting the DTI landscape

One of the advantages of the co-embedding approach that our model takes is the ability to visualize and interpret the shared embedding space, and to use proximity within this space to make additional predictions beyond DTI. We apply a ConPLex model trained on BindingDB and fine-tuned on DUD-E to co-embed cell surface proteins from the Surfaceome database (17). In Figure 4a, we show the projections of 2716 proteins from Surfaceome, colored by their classification into one of five functional categories (from Almén et al. (37)) – transporters, receptors, enzymes, miscellaneous, and those that are unclassified. ConPLex projections of surface proteins cluster in embedding space by functional type, with transporters and receptors especially separating from other classes.

**Fig. 4.**
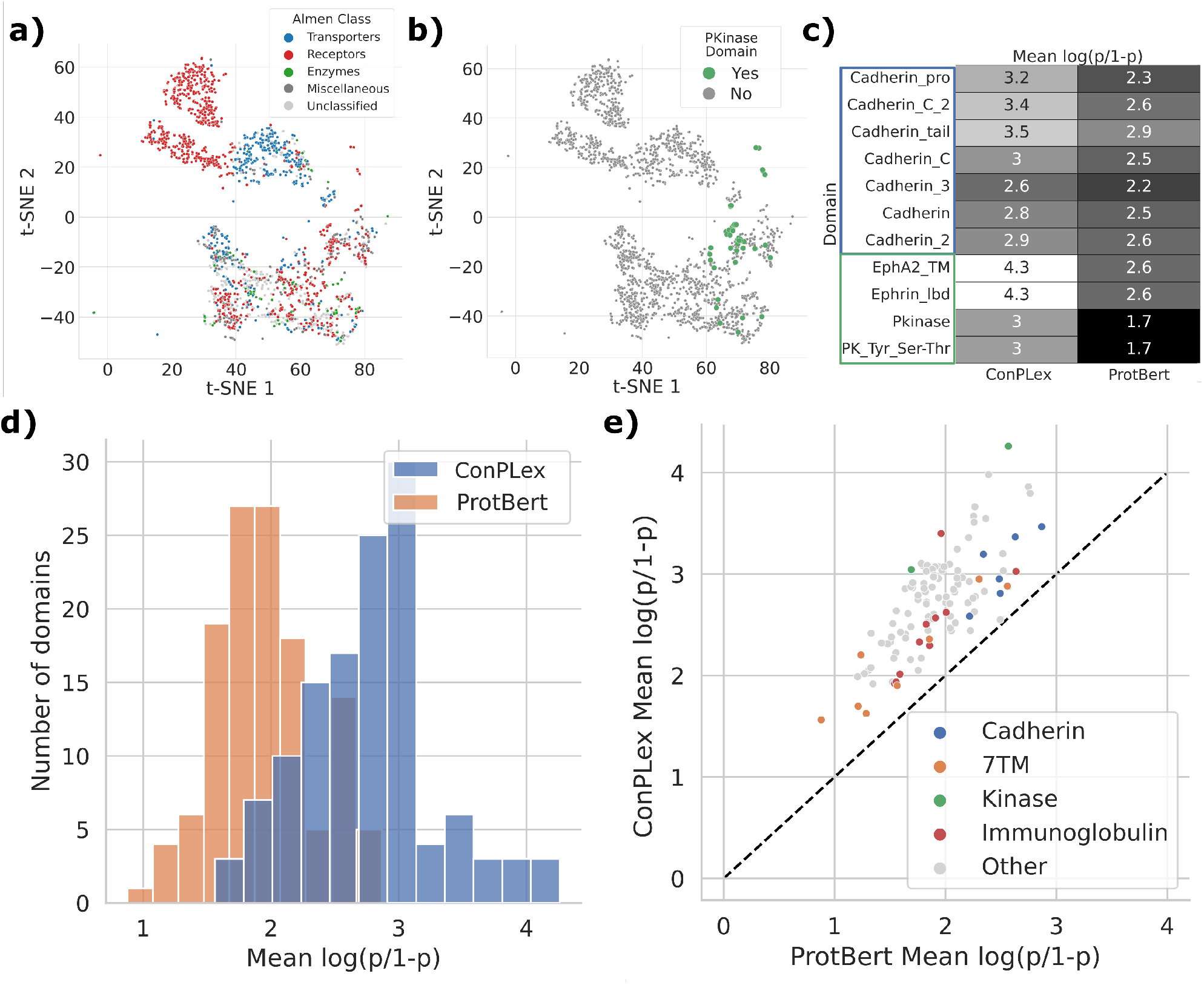
The shared representation space learned by ConPLex is interpretable and captures protein function. **(a)** ConPLex representations of cell surface proteins from the Surfaceome (17) cluster by functional class as assigned in Almén et al. (37). **(b)** These representations also cluster by several functional Pfam domains (38), like the PKinase domain (PF00069) shown in green. **(c)** We evaluated the coherence of representations for each domain by training a logistic regression classifier and report the model’s average confidence for proteins containing that domain as 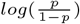. While ConPLex separates all domains better than the untransformed ProtBert embeddings, it differentiates kinase domains (green) especially well, while doing more poorly on cadherin domains (blue). **(d)** Distribution of domain separation for all 126 domains represented in ≥ 10 proteins (*p* = 4.85 × 10^−54^, paired t-test). **(e)** Domain separation using ProtBert vs. ConPLex representations for all 126 domains. ConPLex improves for every domain. We have highlighted several classes of domains, including cadherins (blue), 7-transmembrane proteins (orange), kinases (green), and immunoglobulins (red).

However, the Almén functional classification is quite broad and may group proteins with vastly different functions and binding properties. We further demonstrate the link between ConPLex projections and protein function, by evaluating how the learned DTI embedding space separates proteins by domains contained therein. We identified Pfam domains (38) for each protein in the Surfaceome database using HMMscan (39) and compared the projections of proteins that share the same domains. We identified 780 unique domains across all proteins, of which 126 domains were represented in at least 10 proteins. To quantitatively evaluate the coherence of ConPLex embeddings, we trained separate logistic regression classifiers for each domain to separate proteins with that domain from others, and used the model’s confidence 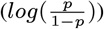 for in-sample proteins as a measure of separation for the domain. We find that for all 126 domains, the model more confidently discriminated domains when trained on ConPLex representations than the baseline ProtBert embeddings.

Figure 4d shows the distributions of all such scores for each model, while Figure 4e shows the change in confidence scores for all 126 domains, where the dotted line represents equal confidence using either ConPLex or ProtBert. We find that while prediction of all domains were improved using ConPLex, kinase domains (PF14575, PF01404, PF00069, PF07714) separated especially well, while Cadherin domains (PF08785, PF16492, PF15974, PF01049, PF16184, PF00028, PF08266) showed more modest improvement (Figure 4c). As discussed previously, ConPLex was trained contrastively with several kinase targets and excels at kinase prediction on DUD-E (Figure 3e), so it is unsuprising that proteins with these domains separate well. In fact, one of the top differentiated domains is the Ephrin ligand binding domain (PF01404), which is responsible for binding to the ephrin ligand (40). In Figure 4b, we show the same visualization of projections as in Figure 4a, but colored by another top-differentiated domain, PKinase (PF00069). While the model was not explicitly trained on targets with Cadherin domains, cadherins typically are responsible for cell-cell adhesion and structural stability, and are less involved in ligand binding (41). Thus, we would not expect a model trained to represent small molecule binding to explicitly differentiate cadherins. This demonstrates that the latent space learned by ConPLex is useful not only for predicting drug-target interaction, but for broadly classifying protein function as it relates to ligand binding. Future work in this area might adjust distances in this landscape to account for the low metric entropy of biological sequences, as demonstrated in Berger, Waterman, and Yu (42).

### Adapting ConPLex for affinity prediction

While we have to this point been using the model to predict probabilities of interaction and perform binary classification, we show that ConPLex can be easily adapted to perform binding affinity prediction, and that this model too achieves state-of-the-art performance. The final step of our binary interaction predictor is converting the cosine distance between the projections in the DTI space to a probability using a sigmoid activation (**Methods**). However, it is completely natural to replace this activation with a dot product between the two projections, which enables the model to make real-valued predictions, which can then be interpreted as a binding affinity. We evaluated ConPLex trained for affinity prediction on the Therapeautics Data Commons (TDC) DTI Domain Generalization (**TDC-DG**) benchmark. The TDC-DG benchmark contains binding affinity (*K*_*d*_) data from interactions patented between 2013 and 2018, with the test set drawn from interactions patented in 2019-2021 (**Methods**). Thus, this data requires out-of-domain generalization and corresponds with the real-life scenario of training on interactions up to a known point, and predicting interactions which are yet to be documented. We trained ConPLex to predict binding affinity with 5 random train/validation splits, and achieve an average Pearson correlation coefficient between the true and predicted affinity of 0.538(±0.008) on the held-out test set. At the time of submission, ConPLex is the top performing method on the TDC-DG benchmark on TDC (Table 2).

**Table 2.** ConPLex can be adapted for state-of-the-art *K*_*d*_ prediction. By replacing the cosine distance in the final step of ConPLex with a dot product between the projections, ConPLex can be used for affinity prediction rather than binary classification. The Therapeatics Data Commons-Domain Generalization (TDC-DG) data set contains *K*_*d*_ values for patented drug-target pairs, where training/testing data are split from before/after 2018. We report the average and standard deviation of the Pearson Correlation Coefficient between true and predicted values across five train/validation splits. Metrics for all methods other than ConPLex come from the TDC leaderboard (19, 43), where at the time of submission ConPLex is the best performing method.

| Model | PCC |
| --- | --- |
| ConPLex | **$0.538 \pm 0.008$** |
| MMD | $0.433 \pm 0.010$ |
| CORAL | $0.432 \pm 0.010$ |
| ERM | $0.427 \pm 0.012$ |
| MTL | $0.425 \pm 0.010$ |
| GroupDRO | $0.384 \pm 0.006$ |
| AndMASK | $0.288 \pm 0.019$ |
| IRM | $0.284 \pm 0.021$ |

## Discussion

Much previous work has recognized the value of meaningful drug representations (44, 45) for DTI prediction, yet relatively little work has focused on the target protein representation. As the first method to use pre-trained protein language models (PLMs) for DTI prediction, ConPLex is yet another example of the power of transferring learned representations for biology (13, 32, 33, 46, 47). This approach enables broad generalization to unseen proteins, as well as extremely fast model inference (*>*10x speed-up even over other sequence-based approaches, Supplementary S4). This speed is particularly valuable for drug re-purposing and iterative screening, where large compound libraries are evaluated against hitherto-uncharacterized proteins implicated in a disease of interest. The co-embedding approach which enables this speedup could also be effective for integrative multi-structure models (e.g., the IMP framework (48)) where efficient scanning of possible combinations is important. Recent methods have also demonstrated the power of PLMs for transferring knowledge between species (33), and our framework may enable more accurate transfer of DTI from the model organisms on which drugs are initially tested, to their eventual use in human patients. Skolnick and Zhou (49) have reported the importance of considering small molecule binding pockets for protein-protein interaction prediction; thus our DTI-informed protein representations may also be useful in that context. While structural similarity is often implicitly learned by PLMs, future work could explicitly incorporate structure where such data is available, perhaps by incorporating a more advanced projection architecture like the Geoformer (3).

It has been shown in previous work that the performance of different PLMs vary on different tasks, and that there is not one clearly “best” language model (14, 50, 51). While we have chosen to use ProtBert here, it is likely that other existing or newly developed language models may yield better performance for certain types of drugs or targets. Likewise, advancements in drug representation may improve performance— the ConPLex framework is flexible to different input features, and it remains important to experiment with different feature choices for the task at hand (Supplementary S2).

ConPLex approaches the DTI decoy problem from the perspective of adversarial machine learning, where the model must act as a discriminator for adversarial examples from the decoy database. This approach is directly enabled by the co-embedding architecture— to compute the triplet distance loss, the protein and drugs must be co-embedded, and the distance between them must be meaningful and simply computed. Such an approach would not be feasible using a model which concatenates features up front, nor for a model which has significant computation defining the probability of interaction after the co-embedding. Thus the shared lexicographic space in which we embed the proteins, targets, and decoys is key. Future work could explore adapting molecular generation methods such as JT-VAE or HierG2G (52, 53) to directly act as a generator for decoys. High-specificity DTI prediction is valuable beyond decoy detection— greater specificity of inference can help improve personalized medicine or the modeling of drug effects against rare variants from under-represented populations.

It is also important to consider the coverage of the problem to select an appropriate method. If enough data is available, for specific enzyme-family prediction tasks we still recommend the use of single-task models (25). To verify individual interactions, energy-based molecular docking will likely be more accurate, although at the cost of being substantially slower (4). Different classes of computational tools for DTI prediction each have varying strengths, and the highest quality predictions can be achieved by leveraging all of these methods together where each is most fit.

Drug discovery is a fundamental task for human health, yet remains both extremely cousre expensive and time-consuming, with the median drug requiring about 1 billion dollars (54) and 10 years (55) from development to approval and distribution. While experimental results will remain the gold-standard for validating drug functionality, *in silico* prediction of drug-target binding remains much faster and cheaper and so will continue to play an important role in early screening of therapeutic candidates (56). To address this step in the drug design pipeline, we have introduced ConPLex. DTI prediction methods should be able to generalize to unseen types of drugs and targets, while also discriminating between highly similar molecules with different binding properties. ConPLex tackles both of these challenges through its dual use of protein language models and contrastive learning. We hope that its broad applicability, specificity on decoys, and ability to scale to massive data will allow ConPLex to be a critical step in this pipeline and contribute to the efficient discovery of effective therapeutics.

## Materials and Methods

### Computing Data Set Coverage

Let 1_(*i,j*)_ be the indicator variable meaning there exists an observation of drug *i* and target *j*. For a data set with *m* unique drugs and *n* unique targets, we can define the coverage for drug *d* as 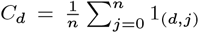 and for a target *t* as 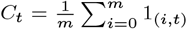. Then, for a given data set we can evaluate the median drug and target coverage. A data set with maximum coverage would have a single data point for each drug-target pair, and thus a median coverage of 1 for both drugs and targets. Conversely, each drug and target would only be represented a single time in a minimum coverage data set, resulting in drug and target coverages of 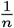 and 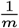 respectively. We report the median drug and target coverage for each benchmark data set in Table 3. Since the DUD-E data set is separated out by targets, we instead report the median number of drugs against each target.

**Table 3.** Full specification of benchmark data sets. We report the number of unique drugs and targets, the median (drug/target) coverage, and the number of training, validation, and test samples in each data set. Number of pairs are shown as (positive/negative), except for TDC-DG (19, 43), which is a regression task, thus total number of pairs is shown. We consider BIOSNAP (57), BindingDB (58), DAVIS (59), and TDC-DG as low-coverage, while Phosphatase (60), Esterase (61), Glycosyltransferase (62), Halogenase (63), BKACE (64), and DUD-E (35) are considered high-coverage. † Because DUD-E is a decoy data set, we report as coverage the median number of true drugs or decoys for each target.

| Data Set | Drugs | Targets | Median Coverage | # Training | # Validation | # Test |
| --- | --- | --- | --- | --- | --- | --- |
| BIOSNAP | 4510 | 2181 | 0.0023 / 0.0020 | 9670 / 9568 | 1396 / 1352 | 2770 / 2727 |
| Unseen Drugs |  |  |  | 9535 / 9616 | 1383 / 1353 | 2918 / 2675 |
| Unseen Targets |  |  |  | 9876 / 9499 | 1382 / 1386 | 2578 / 2762 |
| BindingDB | 7165 | 1254 | 0.0008 / 0.0010 | 6334 / 6334 | 927 / 5717 | 1905 / 11384 |
| DAVIS | 68 | 379 | 0.3707 / 0.3676 | 1043 / 1043 | 160 / 2846 | 303 / 5708 |
| TDC-DG | 140746 | 477 | 0.0021 / 0.0005 | 146891 | 36539 | 49028 |
| Phosphatase | 165 | 218 | 1.0 / 1.0 | 5054 / 27286 | — | 370 / 3260 |
| Esterase | 96 | 146 | 1.0 / 1.0 | 2150 / 10426 | — | 926 / 514 |
| Glycosyltransferase | 89 | 54 | 0.9259 / 0.9778 | 725 / 3042 | — | 113 / 417 |
| Halogenase | 62 | 42 | 1.0 / 1.0 | 303 / 1991 | — | 20 / 290 |
| BKACE | 17 | 161 | 1.0 / 1.0 | 255 / 2193 | — | 19 / 270 |
| DUD-E † |  |  |  | 8996 / 406208 | — | 11430 / 521132 |
| GPCR | 99671 | 5 | 18563 |  |  |  |
| Kinase | 315399 | 26 | 15409 |  |  |  |
| Protease | 286089 | 15 | 9271 |  |  |  |
| Nuclear | 151133 | 11 | 16257 |  |  |  |

### Benchmarks Overview

#### Low coverage benchmarks

We evaluate our framework on three broad-scale, low-coverage benchmark data sets. Two data sets, **DAVIS** (59) and **BindingDB** (58), consist of pairs of drugs and targets with experimentally determined dissociation constants (*K*_*d*_). Following (13), we treat pairs with *K*_*d*_ *<* 30 as positive DTIs, while larger *K*_*d*_ values are negative. The third data set, ChG-Miner from **BIOSNAP** (57), consists of only positive DTIs. We create negative DTIs by randomly sampling an equal number of protein-drug pairs, making the assumption that a random pair is unlikely to be positively interacting. The DAVIS data set represents a few-shot learning setting: it contains only 2,086 training interactions, compared to 12,668 for BindingDB and 19,238 for BIOSNAP. The rest of the data preparation follows (13). The data sets are split into 70% for training, 10% for validation, and the remaining 20% for testing. Training data are artificially sub-sampled to have an equal number of positive and negative interactions, while validation and test data is left at the ratio originally in the data set.

#### Zero-shot benchmarks

We evaluate our framework on two zero-shot prediction modifications of BIOSNAP. Following (13), the **Unseen proteins** set was created by selecting 20% of proteins from the full set, and selecting any interactions including these proteins for the test set. Thus, there are no proteins which appear in both the training and test set. The corresponding process was used to create the **Unseen drugs** data set. The training set was then further split using 7*/*8 of the interactions for training and 1*/*8 of the interactions for testing. As above, data are sub-sampled so that training is balanced.

#### Continuous benchmarks

Continuous affinity prediction data come from the Therapeutics Data Commons DTI Domain Generalization benchmark (**TDC-DG**) (19). The TDC-DG consists of 140,746 unique drugs and 477 unique targets derived from BindingDB (58) interactions that have patent information. Each interaction is labeled with an experimentally determined dissociation constant (*K*_*d*_). Interactions are temporally split, so that training pairs are from patents filed between 2013 and 2018, and test pairs are from between 2019 and 2021. 20% of the training pairs are randomly set aside as a validation set. We train 5 different models with the train/validation splits determined by the TDC benchmarking framework, and report the average Pearson correlation coefficient of predictions on the test set.

#### High coverage benchmarks

The Database of Useful Decoys: Enhanced (**DUD-E**) (35) consists of 102 protein targets and known binding partners (average 224 molecules per target). For each binding partner, there are 50 “decoys”, or physio-chemically similar compounds that are known not to bind with the target. 57 of the targets are classified as either GPCRs, kinases, nuclear proteins, or proteases. We generate train-test splits by splitting targets within classes, so that there are representative members of each class in both the training and test set, but no target appears in both the training and test set (26 train, 31 test). These data are by definition high-coverage, since there are several true and decoy compounds available for each target. We provide the full target splits in Supplementary S1.

We also evaluate on several protein-family specific data sets from various different sources and compiled by Goldman et al. (25). These include DTI data on *β*-ketoacid cleavage (**BKACE**) (64), **Esterase** (61), **Glycosyltransferases** (62), **Halogenase** (63), and **Phosphatase** (60) enzymes. These data are uniformly very high coverage, with a known data point for nearly every drug-target pair. Following (25), we performed a 10-fold cross validation where the data were split into train-test sets by target, so that all drugs appear in both the training and test set, but no target does.

### ConPLex Model

#### Target featurization

We generate protein target features using pretrained protein language models (PLM): These models generate a protein embedding 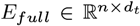 for a protein of length *n*, which is then mean-pooled along the length of the protein resulting in a vector 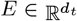. Specifically, we investigate the pre-trained models Prose (31), ESM (65), and ProtBert (27), with default dimensions *d*_*t*_ = 6165, 1280, 1024 respectively (Supplementary S2). Elnaggar et al. recommend the use of ProtT5XLUniref50, but we found that it did not perform as well as ProtBert for the DTI prediction task. We emphasize that the language and projection models are used exclusively to generate input features– their weights are kept unchanged and are not updated during DTI training.

#### Drug featurization

We featurize the drug molecule by its Morgan fingerprint (26), an encoding of the SMILES string of the molecular graph as a fixed-dimension embedding 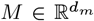 (we chose *d*_*m*_ = 2048) by considering the local neighborhood around each atom. The utility of the Morgan fingerprint for small molecule representation has been demonstrated in (25, 66). We additionally investigated the use of molecule embeddings from Mol2Vec (67) and MolR (68) and found they failed to perform as well as the Morgan fingerprint (Supplementary S2).

#### Transformation into a shared latent space and prediction

Given a target embedding 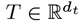 and small molecule embedding 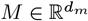, we transform them separately into *T* *, *M* * ∈ ℝ_*h*_ using a single fully-connected layer with a ReLU activation. These layers are parameterized with weight matrices 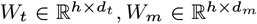 and bias vectors *b*_*t*_, *b*_*m*_ ∈ ℝ^*h*^.

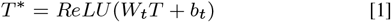

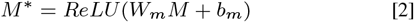

Given the latent embeddings *T* *, *M* *, we compute the probability of a drug-target interaction 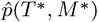 as the cosine similarity between the embedding vectors, followed by a sigmoid activation. Thus, we compute the predicted probability as

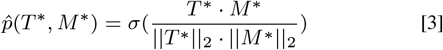

When predicting compound binding affinity *ŷ*(*T*\*, *M*\*), we substitute the sigmoid and cosine similarity (Equation 3) with a dot product followed by a ReLU activation, which gives a non-negative distance in the embedding space (Equation 4).

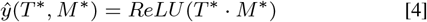

#### Training

The model is trained both for broad and fine prediction, with the loss computed depending on the training data set. Broad-scale training data uses the binary cross-entropy loss (*L*_*BCE*_) between the true labels *y* and the predicted interaction probabilities 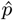. When the model was trained to predict binding affinity, we substitute the binary crossentropy loss for the mean squared error loss (*L*_*MSE*_) is used during supervision.

Training on fine-scale data (DUD-E) was performed using contrastive learning. Contrastive learning uses triplets of training points rather than pairs, denoted the **anchor, positive**, and **negative**, and aims to minimize the distance between the anchor and positive examples while maximizing the distance between the anchor and the negative examples. In the DTI setting, the natural choice for a triplet is the protein target as the anchor, the true drug as the positive and decoy as the negative example, respectively. We derive a training set of triplets in the following manner: for each known interacting drug-target pair (*T, M*^+^), we randomly sample *k* = 50 non-interacting pairs (*T, M*^−^) and generate the triplets (*T, M*^+^, *M*^−^), where *M*^−^ is drawn from the set of all decoys against *T*. We map these to latent space embeddings as described above. Since all the entities are now comparable to each other, we can compute the triplet margin-distance loss (*L*_*TRM*_).

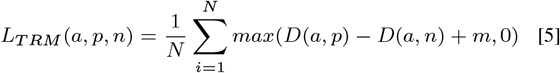

Where

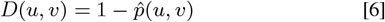

The margin *m* sets the maximum required delta between distances, above which the loss is zero.

#### Margin annealing

The margin *m* sets the maximum required delta between distances, above which the loss is zero. Initially, a large margin requires the decoy to be much further from the target than the drug to avoid a penalty, resulting in larger weight updates. As training progresses, lower margins relax this constraint, requiring only that the drug be closer than the decoy as *m* → 0. Here, the margin is initialized at *M*_*max*_ = 0.25 according to a tanh decay with restarts decay schedule. Every *E*_*max*_ = 10 contrastive epochs, the margin is reset to the initial *M*_*max*_, for a total of 50 epochs. At epoch *i*, the margin is set to

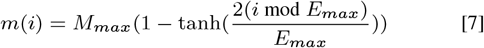

#### Implementation

Model weights were initialized using the Xavier method from a normal distribution (69). Weights were updated with error back-propagation using the AdamW optimizer (70) for a total of 50 epochs. For the binary classification task, the learning rate was initially set to 10^−4^ and adjusted according to a cosine annealing schedule with warm restarts (71) every 10 epochs. For the contrastive task, the learning rate was initially set to 10^−5^ and the same annealing schedule was followed. The margin for the contrastive loss was initially set to 0.25 and decreased to a minimum of 0 over 50 epochs according to a tanh decay schedule with restarts every 10 epochs. We used a latent dimension *d* = 1024 (results were robust to even with lower dimensions, and much higher dimensions may over-fit or be subject to topological restrictions) and a batch size of 32. The model was implemented in PyTorch version 1.11. Model training and inference was performed on a single NVIDIA A100 GPU.

### Surfaceome analysis

We evaluate the interpretability and functional use of ConPLex embeddings using data from the Surfaceome database (17), which contains 2,886 cell-surface proteins. We identified Pfam domains using HMMscan from HMMER3 (39) with default settings. We analyzed domains hit in *>* 10 proteins. For each domain, we trained a logistic regression classifier from sklearn with balanced class weights. We also evaluated domain coherence using spectral clustering with *k* = 10 clusters, and evaluated the adjusted mutual information (AMI) between true clusters (protein has/doesn’t have domain) and predicted clusters (Supplementary S6).

### Genome wide ChEMBL scan

We trained a ConPLex model using BindingDB and DUD-E, and used it to make predictions for all pairs of human proteins against all drugs in ChEMBL. Human protein sequences were taken from the STRING database and processed following (33), resulting in 15,816 proteins between 50 and 800 amino acids long. Small molecule structures were downloaded from ChEMBL 30 (18), resulting in 1,533,652 compounds. Prediction took just under a day, accounting for embedding time.

## ACKNOWLEDGMENTS

RS and BB were supported by the NIH grant R35GM141861. SS was supported by the National Science Foundation Graduate Research Fellowship under Grant No. 2141064. LC was supported by CCF-1934553. The authors thank Kapil Devkota, Tristan Bepler, and Tim Truong for helpful discussions.

## References

1. Jumper J, et al. (2021) Highly accurate protein structure prediction with AlphaFold. Nature 596(7873):583–589.

2. Baek M, et al. (2021) Accurate prediction of protein structures and interactions using a three-track neural network. Science 373(6557):871–876.

3. Wu R, et al. (2022) High-resolution de novo structure prediction from primary se-quence. bioRxiv.

4. Pinzi L, Rastelli G (2019) Molecular docking: shifting paradigms in drug discovery. International journal of molecular sciences 20(18):4331.

5. Bonk BM, Tarasova Y, Hicks MA, Tidor B, Prather KL (2018) Rational design of thiolase substrate specificity for metabolic engineering applications. Biotechnology and bioengineering 115(9):2167–2182.

6. de Melo-Minardi RC, Bastard K, Artiguenave F (2010) Identification of subfamily-specific sites based on active sites modeling and clustering. Bioinformatics 26(24):3075–3082.

7. Trudeau SJ, et al. (2022) Prepci: A structure-and chemical similarity-informed database of predicted protein compound interactions. bioRxiv.

8. Singh R, Park D, Xu J, Hosur R, Berger B (2010) Struct2Net: a web service to predict protein–protein interactions using a structure-based approach. Nucleic acids research 38(suppl_2):W508–W515. publisher: Oxford University Press.

9. Anderson E, Veith GD, Weininger D (1987) SMILES, a line notation and computerized interpreter for chemical structures. (US Environmental Protection Agency, Environmental Research Laboratory).

10. Bagherian M, et al. (2021) Machine learning approaches and databases for prediction of drug–target interaction: a survey paper. Briefings in Bioinformatics p. 23.

11. Hie B, Cho H, Berger B (2018) Realizing private and practical pharmacological collaboration. Science 362(6412):347–350.

12. Lee I, Keum J, Nam H (2019) DeepConv-DTI: Prediction of drug-target interactions via deep learning with convolution on protein sequences. PLoS computational biology 15(6):e1007129.

13. Huang K, Xiao C, Glass LM, Sun J (2021) MolTrans: Molecular Interaction Transformer for drug–target interaction prediction. Bioinformatics 37(6):830–836.

14. Sledzieski S, Singh R, Cowen L, Berger B (2021) Adapting protein language models for rapid DTI prediction. Machine Learning for Structural Biology Workshop (MLSB) at NeurIPS.

15. Bommasani R, et al. (2021) On the opportunities and risks of foundation models. arXiv preprint arXiv:2108.07258.

16. Gururangan S, et al. (2020) Don’t stop pretraining: adapt language models to domains and tasks. arXiv preprint arXiv:2004.10964.

17. Bausch-Fluck D, et al. (2018) The in silico human surfaceome. Proceedings of the National Academy of Sciences 115(46):E10988–E10997.

18. Mendez D, et al. (2019) ChEMBL: towards direct deposition of bioassay data. Nucleic acids research 47(D1):D930–D940.

19. Huang K, et al. (2021) Therapeutics data commons: Machine learning datasets and tasks for drug discovery and development. arXiv preprint arXiv:2102.09548.

20. Zong N, et al. (2022) Beta: a comprehensive benchmark for computational drug– target prediction. Briefings in Bioinformatics.

21. Dönertaş HM, Fuentealba Valenzuela M, Partridge L, Thornton JM (2018) Gene expression-based drug repurposing to target aging. Aging cell 17(5):e12819.

22. Morselli Gysi D, et al. (2021) Network medicine framework for identifying drug-repurposing opportunities for covid-19. Proceedings of the National Academy of Sciences 118(19):e2025581118.

23. Huang K, et al. (2020) DeepPurpose: a deep learning library for drug–target interaction prediction. Bioinformatics 36(22-23):5545–5547.

24. Huang Y, et al. (2019) A framework for identification of on-and off-target transcriptional responses to drug treatment. Scientific reports 9(1):1–9.

25. Goldman S, Das R, Yang KK, Coley CW (2022) Machine learning modeling of family wide enzyme-substrate specificity screens. PLoS computational biology 18(2):e1009853.

26. Morgan HL (1965) The generation of a unique machine description for chemical structures-a technique developed at chemical abstracts service. Journal of Chemical Documentation 5(2):107–113.

27. Elnaggar A, et al. (2020) ProtTrans: towards cracking the language of life’s code through self-supervised deep learning and high performance computing. arXiv preprint arXiv:2007.06225.

28. Tsubaki M, Tomii K, Sese J (2019) Compound–protein interaction prediction with end-to-end learning of neural networks for graphs and sequences. Bioinformatics 35(2):309–318.

29. Scaiewicz A, Levitt M (2015) The language of the protein universe. Current opinion in genetics & development 35:50–56.

30. Bepler T, Berger B (2019) Learning protein sequence embeddings using information from structure in 7th International Conference on Learning Representations, ICLR 2019.

31. Bepler T, Berger B (2021) Learning the protein language: Evolution, structure, and function. Cell Systems 12(6):654–669.e3. Publisher: Elsevier.

32. Heinzinger M, et al. (2019) Modeling aspects of the language of life through transferlearning protein sequences. BMC bioinformatics 20(1):1–17.

33. Sledzieski S, Singh R, Cowen L, Berger B (2021) D-SCRIPT translates genome to phenome with sequence-based, structure-aware, genome-scale predictions of protein-protein interactions. Cell Systems 12:1–14.

34. Singh R, Devkota K, Sledzieski S, Berger B, Cowen L (2022) Topsy-Turvy: integrating a global view into sequence-based ppi prediction. Bioinformatics 38(Supplement_1):i264–i272.

35. Mysinger MM, Carchia M, Irwin JJ, Shoichet BK (2012) Directory of useful decoys, enhanced (DUD-E): better ligands and decoys for better benchmarking. Journal of medicinal chemistry 55(14):6582–6594.

36. Heinzinger M, et al. (2022) Contrastive learning on protein embeddings enlightens midnight zone. NAR genomics and bioinformatics 4(2):lqac043.

37. Almén MS, Nordström KJ, Fredriksson R, Schiöth HB (2009) Mapping the human membrane proteome: a majority of the human membrane proteins can be classified according to function and evolutionary origin. BMC biology 7(1):1–14.

38. El-Gebali S, et al. (2019) The Pfam protein families database in 2019. Nucleic Acids Research 47(D1):D427–D432. Publisher: Oxford University Press.

39. Eddy SR (2011) Accelerated profile HMM searches. PLoS computational biology 7(10):e1002195.

40. Himanen JP, Henkemeyer M, Nikolov DB (1998) Crystal structure of the ligand-binding domain of the receptor tyrosine kinase ephb2. Nature 396(6710):486–491.

41. Maître JL, Heisenberg CP (2013) Three functions of cadherins in cell adhesion. Current Biology 23(14):R626–R633.

42. Berger B, Waterman MS, Yu YW (2020) Levenshtein distance, sequence comparison and biological database search. IEEE transactions on information theory 67(6):3287–3294.

43. Gulrajani I, Lopez-Paz D (2020) In search of lost domain generalization. arXiv preprint arXiv:2007.01434.

44. Huang K, et al. (2021) DeepPurpose: a deep learning library for drug–target interaction prediction. Bioinformatics 36(22-23):5545–5547.

45. Ramsundar B (2018) Ph.D. thesis (Stanford University).

46. Hie B, Zhong ED, Berger B, Bryson B (2021) Learning the language of viral evolution and escape. Science 371(6526):284–288.

47. Littmann M, Heinzinger M, Dallago C, Weissenow K, Rost B (2021) Protein embeddings and deep learning predict binding residues for various ligand classes. Scientific Reports 11(1):1–15.

48. Russel D, et al. (2012) Putting the pieces together: integrative modeling platform software for structure determination of macromolecular assemblies. PLoS biology 10(1):e1001244.

49. Skolnick J, Zhou H (2022) Implications of the essential role of small molecule ligand binding pockets in protein–protein interactions. The Journal of Physical Chemistry B 126(36):6853–6867.

50. Hie BL, Yang KK, Kim PS (2021) Evolutionary velocity with protein language models. bioRxiv.

51. Hsu C, Nisonoff H, Fannjiang C, Listgarten J (2021) Combining evolutionary and assay-labelled data for protein fitness prediction. bioRxiv.

52. Jin W, Barzilay R, Jaakkola T (2018) Junction tree variational autoencoder for molecular graph generation in International Conference on Machine Learning. (PMLR), pp. 2323–2332.

53. Jin W, Barzilay R, Jaakkola T (2020) Hierarchical generation of molecular graphs using structural motifs in International Conference on Machine Learning. (PMLR), pp. 4839–4848.

54. Wouters OJ, McKee M, Luyten J (2020) Estimated research and development investment needed to bring a new medicine to market, 2009-2018. Jama 323(9):844–853.

55. Van Norman GA (2016) Drugs, devices, and the fda: part 1: an overview of approval processes for drugs. JACC: Basic to Translational Science 1(3):170–179.

56. Sabe VT, et al. (2021) Current trends in computer aided drug design and a highlight of drugs discovered via computational techniques: A review. European Journal of Medicinal Chemistry 224:113705.

57. Zitnik M, Sosič R, Maheshwari S, Leskovec J (2018) BioSNAP Datasets: Stanford biomedical network dataset collection (http://snap.stanford.edu/biodata).

58. Liu T, Lin Y, Wen X, Jorissen RN, Gilson MK (2007) BindingDB: a web-accessible database of experimentally determined protein–ligand binding affinities. Nucleic acids research 35(suppl_1):D198–D201.

59. Davis MI, et al. (2011) Comprehensive analysis of kinase inhibitor selectivity. Nature biotechnology 29(11):1046–1051.

60. Huang H, et al. (2015) Panoramic view of a superfamily of phosphatases through substrate profiling. Proceedings of the National Academy of Sciences 112(16):E1974–E1983.

61. Martínez-Martínez M, et al. (2017) Determinants and prediction of esterase substrate promiscuity patterns. ACS chemical biology 13(1):225–234.

62. Yang M, et al. (2018) Functional and informatics analysis enables glycosyltransferase activity prediction. Nature chemical biology 14(12):1109–1117.

63. Fisher BF, Snodgrass HM, Jones KA, Andorfer MC, Lewis JC (2019) Site-selective c– h halogenation using flavin-dependent halogenases identified via family-wide activity profiling. ACS central science 5(11):1844–1856.

64. Bastard K, et al. (2014) Revealing the hidden functional diversity of an enzyme family. Nature chemical biology 10(1):42–49.

65. Rives A, et al. (2021) Biological structure and function emerge from scaling unsupervised learning to 250 million protein sequences. Proceedings of the National Academy of Sciences 118(15).

66. Rogers D, Hahn M (2010) Extended-connectivity fingerprints. Journal of chemical information and modeling 50(5):742–754.

67. Jaeger S, Fulle S, Turk S (2018) Mol2vec: unsupervised machine learning approach with chemical intuition. Journal of chemical information and modeling 58(1):27–35.

68. Wang H, et al. (2021) Chemical-reaction-aware molecule representation learning. arXiv preprint arXiv:2109.09888.

69. Glorot X, Bengio Y (2010) Understanding the difficulty of training deep feedforward neural networks in Proceedings of the thirteenth international conference on artificial intelligence and statistics. (JMLR Workshop and Conference Proceedings), pp. 249–256.

70. Loshchilov I, Hutter F (2017) Decoupled weight decay regularization. arXiv preprint arXiv:1711.05101.

71. Loshchilov I, Hutter F (2016) Sgdr: Stochastic gradient descent with warm restarts. arXiv preprint arXiv:1608.03983.

