## Supplemental Information for "Learning the Drug-Target Interaction Lexicon"

#### This PDF file includes:

Supporting text
Figs. S1 to S2
Tables S1 to S6
SI References

### Supporting Information Text

### 1. DUD-E Train/Test Splits

Targets were randomly split within each class, so that there were examples of each class in the training and test set. There are a total of 57 targets, 26 in training and 31 in testing. A list of all targets and classes evaluated in train and test are in Table S1.

**Table S1. DUD-E target classes and splits.**

| Target ID | Class | Split |
|---|---|---|
| AA2AR | Gpcr | Train |
| ABL1 | Kinase | Train |
| AKT1 | Kinase | Train |
| CDK2 | Kinase | Train |
| EGFR | Kinase | Train |
| FAK1 | Kinase | Train |
| FGFR1 | Kinase | Train |
| IGF1R | Kinase | Train |
| JAK2 | Kinase | Train |
| KIT | Kinase | Train |
| MAPK2 | Kinase | Train |
| MK01 | Kinase | Train |
| PLK1 | Kinase | Train |
| TGFR1 | Kinase | Train |
| ANDR | Nuclear | Train |
| ESR2 | Nuclear | Train |
| MCR | Nuclear | Train |
| PPARD | Nuclear | Train |
| THB | Nuclear | Train |
| ACE | Protease | Train |
| BACE1 | Protease | Train |
| DPP4 | Protease | Train |
| FA7 | Protease | Train |
| HIVPR | Protease | Train |
| MMP13 | Protease | Train |
| TRYB1 | Protease | Train |
| ADRB1 | Gpcr | Test |
| ADRB2 | Gpcr | Test |
| CXCR4 | Gpcr | Test |
| DRD3 | Gpcr | Test |
| AKT2 | Kinase | Test |
| BRAF | Kinase | Test |
| CSF1R | Kinase | Test |
| KPCB | Kinase | Test |
| LCK | Kinase | Test |
| MET | Kinase | Test |
| MK10 | Kinase | Test |
| MK14 | Kinase | Test |
| MP2K1 | Kinase | Test |
| ROCK1 | Kinase | Test |
| SRC | Kinase | Test |
| VGFR2 | Kinase | Test |
| WEE1 | Kinase | Test |
| ESR1 | Nuclear | Test |
| GCR | Nuclear | Test |
| PPARA | Nuclear | Test |
| PPARG | Nuclear | Test |
| PRGR | Nuclear | Test |
| RXRA | Nuclear | Test |
| ADA17 | Protease | Test |
| CASP3 | Protease | Test |
| FA10 | Protease | Test |
| LKHA4 | Protease | Test |
| RENI | Protease | Test |
| THRB | Protease | Test |
| TRY1 | Protease | Test |
| UROK | Protease | Test |

### 2. Choice of protein and molecule featurization

**A. Alternate language model options for target feature generation.** We explored several choices for using language models to generate protein features, including D-SCRIPT (1), Prose (2, 3), ESM (4), and ProtBert (5). For D-SCRIPT, we use the 100 dimensional embeddings after the first projection layer. For the other language models, we use the output of the final embedding layer. All models provide per-amino acid features, which we average along the length of the protein to get a fixed length vector. Table S2 shows the performance of a ConPLex model using each of the featurizations (without contrastive learning for simplicity). While there is no one language model which is uniformly best (**Discussion**), we chose to use ProtBert which seemed to have consistent strong performance.

**B. Alternate options for drug feature generation.** We also explored three different choices for generating drug features, including the Morgan fingerprint (6), MolR (7), and Mol2Vec (8). Results are shown in Table S3. The Morgan fingerprint uniformly outperforms the other choices, and so we chose to use it for ConPLex .

**C. Feature attribution reveals information gain from tuning on PPI.** We additionally investigated training ConPLex models with features generated by concatenating protein language model embeddings with embeddings from a D-SCRIPT (1) model pre-trained on human PPIs. We refer to the output of the first projection module of D-SCRIPT, which takes as input the $n \times d$ protein representation and tunes it to an $n \times 100$ representation. We then concatenate this with the original PLM representation to get an embedding with $d + 100$ features.

 Rohit Singh*, Samuel Sledzieski*, Lenore Cowen, and Bonnie Berger

**Table S2. ConPLex classification results on benchmark data sets using different PLMs to generate protein features.**

| Benchmark | Protein Features | AUPR | AUROC | F1 |
|---|---|---|---|---|
| BIOSNAP | D-SCRIPT | 0.911 | 0.897 | 0.831 |
| | Prose | **0.914** | **0.898** | **0.838** |
| | ESM | 0.898 | 0.876 | 0.817 |
| | ProtBert | 0.895 | 0.873 | 0.811 |
| BindingDB | D-SCRIPT | 0.618 | 0.870 | 0.619 |
| | Prose | 0.618 | 0.862 | 0.625 |
| | ESM | 0.637 | **0.881** | **0.637** |
| | ProtBert | **0.652** | 0.876 | 0.636 |
| DAVIS | D-SCRIPT | 0.475 | 0.912 | 0.533 |
| | Prose | 0.463 | 0.907 | 0.523 |
| | ESM | 0.480 | 0.916 | 0.544 |
| | ProtBert | **0.511** | **0.917** | **0.546** |
| BIOSNAP Unseen Proteins | Prose | **0.875** | **0.868** | **0.798** |
| | ESM | 0.850 | 0.839 | 0.770 |
| | ProtBert | 0.841 | 0.827 | 0.750 |
| BIOSNAP Unseen Drugs | Prose | **0.881** | **0.857** | **0.802** |
| | ESM | 0.876 | 0.850 | 0.796 |
| | ProtBert | 0.876 | 0.848 | 0.785 |

**Table S3. ConPLex classification results on benchmark data sets using different methods to generate small molecule features.**

| Benchmark | Drug Features | AUPR |
|---|---|---|
| BIOSNAP | Morgan | **0.904** |
| | MolR | 0.853 |
| | Mol2Vec | 0.792 |
| BindingDB | Morgan | **0.668** |
| | MolR | 0.578 |
| | Mol2Vec | 0.380 |
| DAVIS | Morgan | **0.511** |
| | MolR | 0.436 |
| | Mol2Vec | 0.144 |

While the top-line performance of the augmented models are similar to the base models, an attribution study using DeepLift (9) shows that the new D-SCRIPT-derived features are disproportionately represented in the set of highly important features (Figure S1). This suggests that tuning on a related task refines the representations from the general protein language models to ones more suited for the specific task, as in (10). This explanation is supported by the fact that the 100-dimensional D-SCRIPT features alone achieved only slightly decreased performance on the DTI task compared to PLM-based models with 10-50x as many parameters (Table S2).

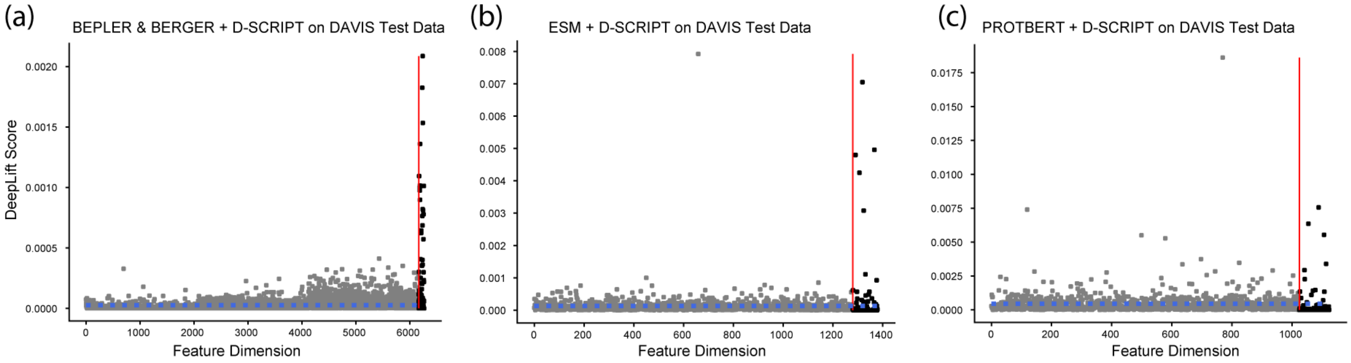


**Fig. S1.** DeepLift feature attributions for Prose **(a)**, ESM **(b)**, and ProtBert **(c)** embedding dimensions (gray) and the respective D-SCRIPT embedding dimensions (black). D-SCRIPT features have an outsize contribution to the overall model prediction relative to the PLM features alone. The dashed blue line indicates the mean feature attribution.

### 3. Margin decay ablation and optimization

We evaluated the impact of various different margin decay schemes, as well as the impact of contrastive learning overall on model performance. We report the average AUPR over 3 random initializations on the BindingDB data set. The baseline without contrastive learning has the highest AUPR, but as we have shown contrastive learning is essential for specificity on fine-scale data sets. The best performing model with contrastive learning is within 5% of the performance without. In addition to the Tanh decay with restart we describe in the main text, we evaluate the performance of a Tanh decay without a restart, a cosine decay with and without restarts, and a constant margin (no decay). We find that the Tanh decay with restart performs the best, and show the full results in Table S4.

**Table S4. Evaluation of different margin decay schemes on BindingDB.**

| Decay Scheme | Restart? | AUPR |
|---:|:---:|:---:|
| **No Contrastive Learning** | N/A | 0.664 |
| Tanh | ✓ | **0.632** |
| Tanh | ✗ | 0.609 |
| Cosine | ✓ | 0.620 |
| Cosine | ✗ | 0.602 |
| Constant | ✗ | 0.595 |

### 4. Training and inference time analysis

One of the benefits of using pre-trained models for feature generation is that computation times are amortized over the lifespan of downstream applications. Pre-trained models incur an up-front computational cost, but can then be re-used for multiple inference tasks with straightforward architectures. Our framework allows for training DTI models up to 8 times faster than an end-to-end method like MolTrans (11). Inference of DTIs is also roughly an order of magnitude faster (Table S5). MolTrans has a faster per-epoch training time than ConPLex on DAVIS, but a slower wall clock time and much slower inference.

**Table S5. Comparison of training and inference times on three high-coverage data sets with contrastive training on DUD-E (mean seconds over 5 runs).**

| Benchmark | Model | Wall Clock | Training (per epoch) | Inference |
|---|---|---:|---:|---:|
| BIOSNAP | ConPLex | **1273** | **22.84** | **0.556** |
| | MolTrans | 6424 | 116.21 | 31.64 |
| BindingDB | ConPLex | **1302** | **24.21** | **2.168** |
| | MolTrans | 9874 | 142.96 | 222.17 |
| DAVIS | ConPLex | **1145** | 22.38 | **0.619** |
| | MolTrans | 1417 | **15.38** | 35.06 |

### 5. Results on EnzPred data sets

Results of ConPLex , EnzPred-CPI, and a single-task ridge regression model on several protein-family specific data sets compiled by Goldman et al. (12). These include $\beta$-ketoacid cleavage (**BKACE**) (13), **Esterase** (14), **Glycosyltransferases** (15), **Halogenase** (16), and **Phosphatase** (17) enzymes. Following (12), we performed a 10-fold cross validation where the data were split into train-test sets by target, so that all drugs appear in both the training and test set, but no target does. We report the average of 3 random initializations in Table S6. "Average AUPR" means AUPR was computed per-drug then averaged.

### 6. Surfaceome analysis with spectral clustering

In addition to measuring the quality of the latent space using a classifier, we also tried an unsupervised approach. We trained a spectral clustering algorithm with $k = 10$ clusters. Then, for each domain, we compute the adjusted mutual information (AMI) between the clustering assignments and the presence/absence of that domain. A higher AMI means that proteins with the same domain are more likely to share a cluster. We find that ConPLex embeddings result in signficantly higher AMIs than ProtBert (paired t-test $p = 5.37e - 4$). We show clustering results for each domain as well as summary histograms in Figure S2. We also show the distribution of domain cluster sizes for all 780 domains identified in at least one protein, and our cutoff that a domain must have $> 10$ proteins for us to consider it in Figure S2D.

**Table S6. Comparison of PLM-based drug-target interaction models with a per-drug Ridge regression model on enzyme-family specificity benchmark data sets.**

| Benchmark | Model | Average AUPR |
|---|---|---|
| BKACE | ConPLex | 0.53 ± 0.04 |
| | EnzPred-CPI | 0.41 ± 0.01 |
| | Ridge (Single-Task) | **0.62** ± **0.02** |
| Esterase | ConPLex | 0.46 ± 0.02 |
| | EnzPred-CPI | 0.52 ± 0.02 |
| | Ridge (Single-Task) | **0.58** ± **0.01** |
| Gylosyltransferase | ConPLex | 0.52 ± 0.03 |
| | EnzPred-CPI | 0.46 ± 0.01 |
| | Ridge (Single-Task) | **0.55** ± 0.00 |
| Halogenase | ConPLex | **0.47** ± **0.04** |
| | EnzPred-CPI | 0.38 ± 0.01 |
| | Ridge (Single-Task) | 0.44 ± 0.03 |
| Phosphatase | ConPLex | 0.33 ± 0.02 |
| | EnzPred-CPI | 0.39 ± 0.01 |
| | Ridge (Single-Task) | **0.40** ± **0.00** |

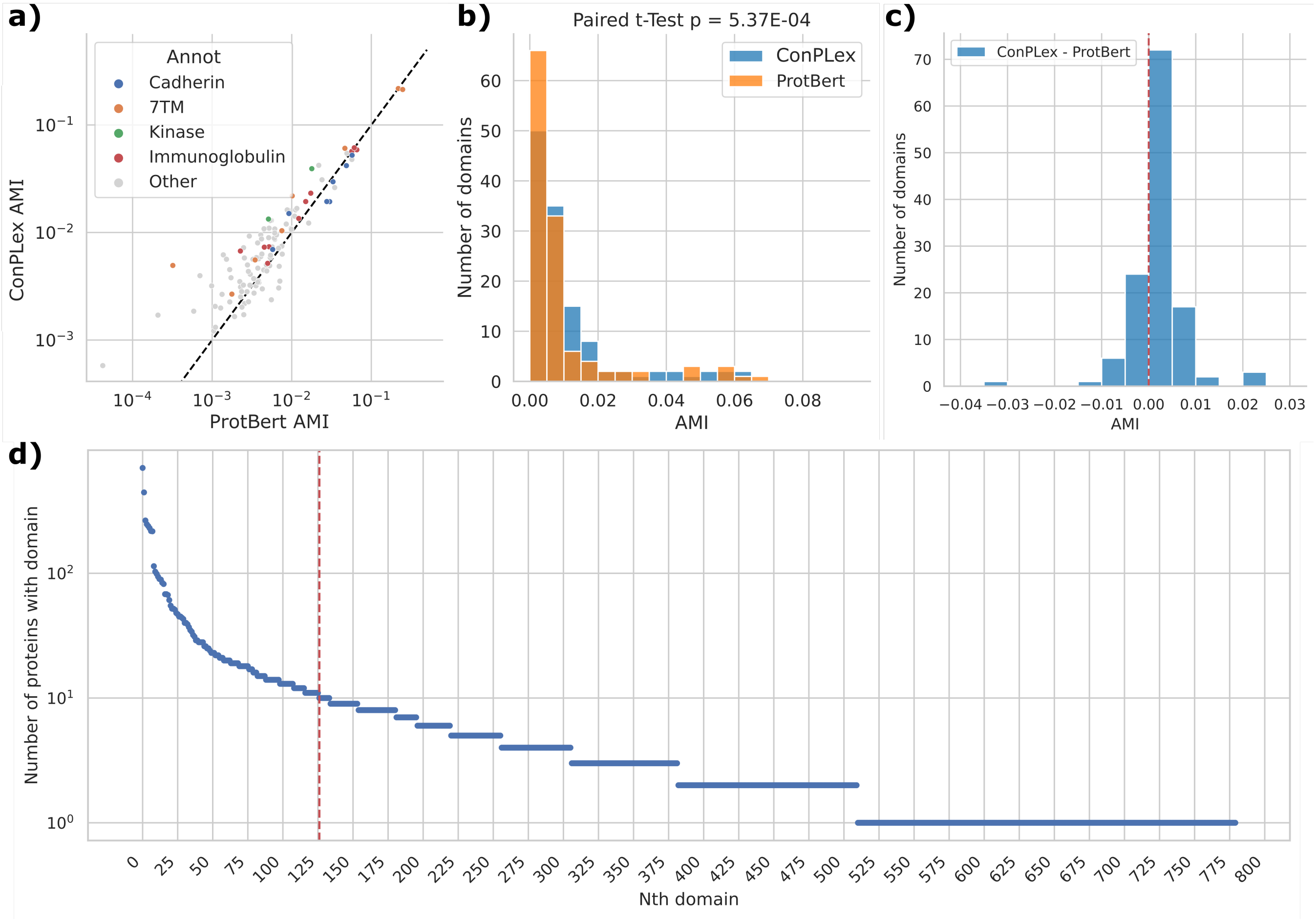


**Fig. S2. Analysis of domain coherence on Surfaceome (a)** ProtBert vs. ConPLex spectral clustering AMI to true separation for each domain. Above the diagonal dotted line means that ConPLex improves upon ProtBert representations. **(b)** Histogram of clustering quality AMI for ConPLex and ProtBert **(c)** Histogram of residuals. To the left of 0, ProtBert performed better, while ConPLex performed better for domains to the right of 0. **(d)** For all domains that were annotated using HMMscan, we plot the number of proteins that have that domain. We don't evaluate any domains that appear in fewer than 10 proteins, since clustering or classification metrics could be subject to noise. As a result, we evaluate on 126 domains.

 Rohit Singh*, Samuel Sledzieski*, Lenore Cowen, and Bonnie Berger

### References


1. Sledzieski S, Singh R, Cowen L, Berger B (2021) D-SCRIPT translates genome to phenome with sequence-based, structure-aware, genome-scale predictions of protein-protein interactions. *Cell Systems* 12:1–14.
2. Bepler T, Berger B (2019) Learning protein sequence embeddings using information from structure in *7th International Conference on Learning Representations, ICLR 2019*.
3. Bepler T, Berger B (2021) Learning the protein language: Evolution, structure, and function. *Cell Systems* 12(6):654–669.e3. Publisher: Elsevier.
4. Rives A, et al. (2021) Biological structure and function emerge from scaling unsupervised learning to 250 million protein sequences. *Proceedings of the National Academy of Sciences* 118(15).
5. Elnaggar A, et al. (2020) ProtTrans: towards cracking the language of life's code through self-supervised deep learning and high performance computing. *arXiv preprint arXiv:2007.06225*.
6. Morgan HL (1965) The generation of a unique machine description for chemical structures-a technique developed at chemical abstracts service. *Journal of Chemical Documentation* 5(2):107–113.
7. Wang H, et al. (2021) Chemical-reaction-aware molecule representation learning. *arXiv preprint arXiv:2109.09888*.
8. Jaeger S, Fulle S, Turk S (2018) Mol2vec: unsupervised machine learning approach with chemical intuition. *Journal of chemical information and modeling* 58(1):27–35.
9. Shrikumar A, Greenside P, Kundaje A (2017) Learning important features through propagating activation differences in *International Conference on Machine Learning*. (PMLR), pp. 3145–3153.
10. Gururangan S, et al. (2020) Don't stop pretraining: adapt language models to domains and tasks. *arXiv preprint arXiv:2004.10964*.
11. Huang K, Xiao C, Glass LM, Sun J (2021) MolTrans: Molecular Interaction Transformer for drug–target interaction prediction. *Bioinformatics* 37(6):830–836.
12. Goldman S, Das R, Yang KK, Coley CW (2022) Machine learning modeling of family wide enzyme-substrate specificity screens. *PLoS computational biology* 18(2):e1009853.
13. Bastard K, et al. (2014) Revealing the hidden functional diversity of an enzyme family. *Nature chemical biology* 10(1):42–49.
14. Martínez-Martínez M, et al. (2017) Determinants and prediction of esterase substrate promiscuity patterns. *ACS chemical biology* 13(1):225–234.
15. Yang M, et al. (2018) Functional and informatics analysis enables glycosyltransferase activity prediction. *Nature chemical biology* 14(12):1109–1117.
16. Fisher BF, Snodgrass HM, Jones KA, Andorfer MC, Lewis JC (2019) Site-selective c–h halogenation using flavin-dependent halogenases identified via family-wide activity profiling. *ACS central science* 5(11):1844–1856.
17. Huang H, et al. (2015) Panoramic view of a superfamily of phosphatases through substrate profiling. *Proceedings of the National Academy of Sciences* 112(16):E1974–E1983.
